# Cyclocreatine suppresses prostate tumorigenesis through dual effects on SAM and creatine metabolism

**DOI:** 10.1101/2021.02.26.432994

**Authors:** Rachana Patel, Lisa Rodgers, Catriona A. Ford, Linda K Rushworth, Janis Fleming, Ernest Mui, Tong Zhang, David Watson, Gillian Mackay, David Sumpton, Owen J. Sansom, Hing Y. Leung

**Affiliations:** CRUK Beatson Institute, Garscube Estate, Switchback Road, Glasgow, G61 1BD, UK; Institue of Cancer Sciences, University of Glasgow, Switchback Road, Glasgow, G61 1QH, UK; Strathclyde Institute of Pharmacy and Biomedical Sciences, University of Strathclyde, 161 Cathedral St, Glasgow, G4 0RE, UK

**Keywords:** Creatine, phosphocreatine, S-adenosyl methionine, cyclocreatine, prostate cancer

## Abstract

Prostate cancer is highly prevalent, being the second most common cause of cancer mortality in men worldwide. Applying a novel genetically engineered mouse model (GEMM) of aggressive prostate cancer driven by deficiency of PTEN and SPRY2 (Sprouty 2) tumour suppressors, we identified enhanced creatine metabolism within the phosphagen system in progressive disease. Altered creatine metabolism was validated in *in vitro* and *in vivo* prostate cancer models and in clinical cases. Upregulated creatine levels were due to increased uptake through the SLC6A8 creatine transporter and *de novo* synthesis, resulting in enhanced cellular basal respiration. Treatment with cyclocreatine (a creatine analogue that potently and specifically blocks the phosphagen system) dramatically reduces creatine and phosphocreatine levels. Blockade of creatine biosynthesis by cyclocreatine leads to cellular accumulation of S-adenosyl methionine (SAM), an intermediary of creatine biosynthesis, and suppresses prostate cancer growth *in vitro*. Furthermore, cyclocreatine treatment impairs cancer progression in our GEMM and in a xenograft liver metastasis model. Hence, by targeting the phosphagen system, cyclocreatine results in anti-tumourigenic effects from both SAM accumulation and suppressed phosphagen system.

## INTRODUCTION

Prostate cancer is the most common cancer in men worldwide (Siegel *et al*, 2020). Around 20% of patients present with advanced disease at the time of diagnosis. Even for men found to have an early or indolent form of the disease, at least 20% of these patients will subsequently progress to a lethal and treatment-resistant form of prostate cancer (Lohiya *et al*, 2016). Therefore, identification and characterisation of genetic lesions that influence aggressive prostate cancer growth may provide new strategies to improve clinical management of disease progression. We have previously shown that genomic loss of the tumour suppressors PTEN and Sprouty2 (SPRY2) co-occur in progressive prostate cancers including in treatment resistance (Gao *et al*, 2012; Patel *et al*, 2018; Patel *et al*, 2013), and that concurrent loss of PTEN and SPRY2 leads to aggressive treatment-resistant disease (Gao *et al*., 2012; Patel *et al*., 2018; Patel *et al*., 2013).

Oncogenic metabolic rewiring mediated by loss of tumour suppressors is one of the hallmarks of cancer (Hanahan & Weinberg, 2011). Identifying, understanding and targeting the metabolic vulnerabilities triggered by loss of tumour suppressors is an appealing approach for designing interventions to tackle cancer growth and invasion. Aggressive cancers have high energy demands and may experience nutrient restricting conditions during rapid proliferative stages of tumorigenesis (Fenouille *et al*, 2017). The phosphagen system refers to high energy storage compounds and associated biosynthesis enzymes (Patra *et al*, 2012; Wallimann, 1994; Wyss & Kaddurah-Daouk, 2000). Cancers of liver and breast exploit the phosphagen system, the cellular energy buffering system, to meet their high energy demands (Qian *et al*, 2012).

The phosphagen system comprises of creatine, a nitrogen amine that can be phosphorylated by creatine kinases to form phosphocreatine. In addition to uptake of creatine from the circulation, creatine can also be synthesised *de novo* from arginine and glycine, with S-adenosyl methionine (SAM) serving as the methyl donor (Bera *et al*, 2008). One of the well-characterised roles of creatine is to act as a cellular energy buffer by rapidly transferring energy through a reversible reaction catalysed by creatine kinases. Creatine kinases play a central role in the energy metabolism of cells that have high and fluctuating energy requirements by catalysing the reversible transfer of the phosphoryl group from phosphocreatine to ADP to generate ATP (Kazak & Cohen, 2020). Phosphocreatine serves as an easily diffusible energy storage metabolite to regenerate ATP on demand by cytosolic creatine kinase isoforms.

The role of creatine metabolism in prostate cancer has not been fully explored. Here, using clinically relevant human and murine prostate cancer models, we have investigated the role of combined PTEN and SPRY2 deficiency in mediating metabolic reprogramming of prostate cancer cells. Our data point to enhanced creatine uptake and biosynthesis in progressive prostate cancer. Importantly, we identified SAM utilisation as an additional mechanism of how cancer cells may use creatine metabolism beyond fuelling cellular bioenergetics to maintain cell growth.

## RESULTS

### Concurrent loss of PTEN and SPRY2 mediates prostate cancer progression

Heterozygous and homozygous genomic deletions of the tumour suppressors *PTEN* and *SPRY2* occur in up to half of metastatic prostate cancers (Figure 1A) (Abida *et al*, 2019; Patel *et al*., 2013). Using a murine model of prostate cancer, we have previously shown that heterozygous loss of *Pten* and *Spry2* cooperate to drive prostate cancer progression and treatment resistance (Patel *et al*., 2018; Patel *et al*., 2013). To investigate the functional co-operation of *Spry2* deficiency with homozygous deletion of *Pten*, we generated a prostate-specific *Pten* null mouse model with heterozygous or homozygous deletion of *Spry2* using a conditional Cre-loxP system driven by *Probasin*-*Cre* (Patel *et al*., 2013). The overall survival of mice with complete deletion of *Pten* and *Spry2* (*Pten*^*pc-/-*^ *Spry2*^*pc-/-*^) was significantly shorter than mice with deletion of *Pten* (*Pten*^*pc-/-*^) alone, deletion of *Spry2* (*Spry2*^*pc-/-*^) alone or *Pten* deletion with heterozygous loss of *Spry2* (*Pten*^*pc-/-*^ *Spry2*^*pc-/+*^) (Figure 1B). As observed before, *Spry2* loss alone did not lead to tumour formation (Patel *et al*., 2013). At the study endpoint, *Pten* null mice with single or double copy loss of *Spry2* had significantly higher prostate tumour burden compared to mice with *Pten* deletion alone (Figure 1C, EV1A). *Pten* and *Spry2* null (*Pten*^*pc-/-*^ *Spry2*^*pc-/-*^) tumours were also highly proliferative with stronger Ki67 staining than the other genotypes (Figure 1D-E).

**Figure 1.**
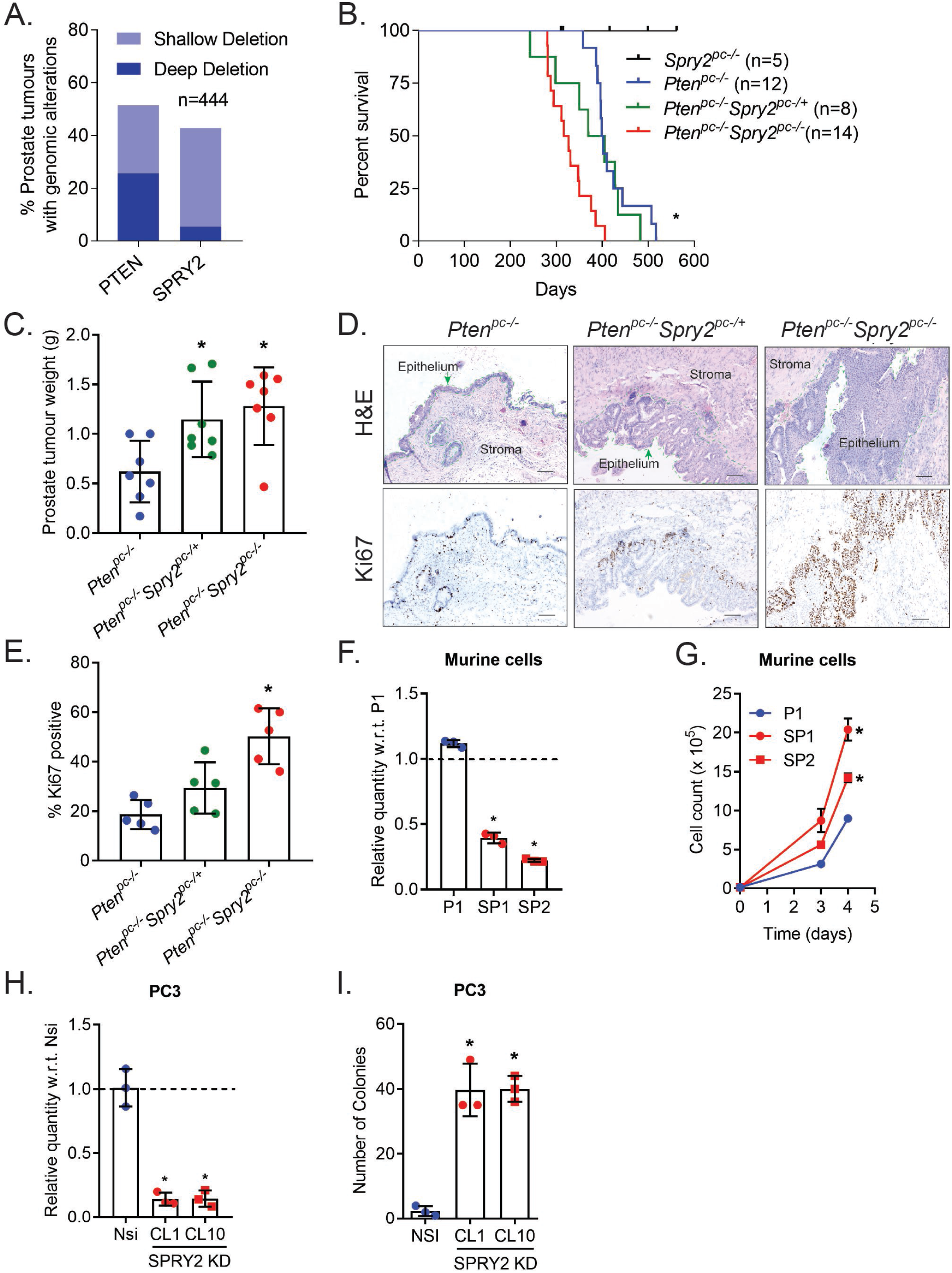
Concomitant PTEN and SPRY2 deficiencies drive prostate cancer progression. *(A) PTEN* and *SPRY2* genomic alterations in metastatic prostate cancer patients (SU2C/PCF Dream Team, PNAS 2019). (B) Kaplan–Meier plot for overall survival of indicated mice (*p<0.05, *Pten*^*pc-/-*^ *Spry2*^*pc-/-*^compared to *Pten*^*pc-/-*^; Log-rank Mantel-Cox test). (C) Non-cystic prostate tumour weights from indicated mice at clinical endpoints (n=7 mice for each group; mean values ± SD are shown; *p<0.05 compared to *Pten*^*pc-/-*^; 1-way ANOVA with Tukey’s multiple comparison test). (D) Representative H&E and Ki67 images of prostate tumour sections from *Pten*^*pc-/-*^, *Pten*^*pc-/-*^ *Spry2*^*pc-/+*^ and *Pten*^*pc-/-*^ *Spry2*^*pc-/-*^ mice (n=5 mice for each group; scale bar = 100 μm). (E) IHC quantification of Ki67 in prostate tumours as indicated (n=5 mice for each group; mean values ± SD are shown; *p<0.05 compared to *Pten*^*pc-/-*^; 1-way ANOVA with Tukey’s multiple comparison test). (F) Relative mRNA levels of *Spry2* in primary murine prostate cancer cells, as indicated. P1 derived from *Pten*^*pc-/-*^; SP1 and SP2 derived from two independent *Pten*^*pc-/-*^ *Spry2*^*pc-/+*^ prostate tumours (n=3 independent experiments; mean values ± SD are shown; *p<0.05 compared to P1; 1-way ANOVA with Dunnett’s multiple comparison test). (G) Growth of indicated cells (n=3 independent experiments; mean values ± SEM are shown; *p<0.05 compared to P1; 2-way ANOVA with Tukey’s multiple comparison test). (H) Relative mRNA levels of *SPRY2* in PC3 human prostate cancer cell lines as indicated. Nsi is PC3 cells with stable expression of non-silencing vector control; CL1 and CL10 are PC3 clones with stable knockdown (KD) of SPRY2 (n=3 independent experiments; mean values ± SD are shown; *p<0.05 compared to Nsi; 1-way ANOVA with Dunnett’s multiple comparison test). (I) Soft agar colony quantifications of the indicated cells (data shown are representative of n=3 independent experiments; mean values ± SD are shown; *p<0.05 compared to Nsi; 1-way ANOVA with Dunnett’s multiple comparison test).

Primary murine prostate cancer SP1 and SP2 cell lines were derived from two independent *Pten*^pc*-/-*^ *Spry2*^*pc-/+*^ double mutant tumours, and confirmed to have suppressed SPRY2 expression (Figure 1F, EV1B). As observed in murine tumours, SP1 and SP2 cells proliferated significantly faster than the P1 cells derived from a *Pten* null tumour (Figure 1G). Thus, concurrent loss of PTEN and SPRY2 tumour suppressors enhances murine prostate tumorigenesis with increased cell proliferation. Similarly, in PTEN deficient human PC3 prostate cancer cells, stable knockdown of SPRY2 expression (CL1 and CL10) significantly promoted their soft agar colony formation efficiency (Figure 1H-I, EV1C).

### PTEN and SPRY2 deficiency leads to increased creatine uptake through SLC6A8

We wished to test if there was evidence of metabolic rewiring in tumours driven by concurrent loss of PTEN and SPRY2. Untargeted analysis of metabolites revealed enriched creatine metabolism in *Pten*^*pc-/-*^ *Spry2*^*pc-/-*^ tumours as well as in SPRY2 deficient PC3 cells (Table S1, Figure EV2A). Targeted metabolomics analysis was then carried out, and significant enrichment of creatine in SPRY2 deficient human and murine prostate cancer cells was confirmed (Figure 2A-B).

**Figure 2.**
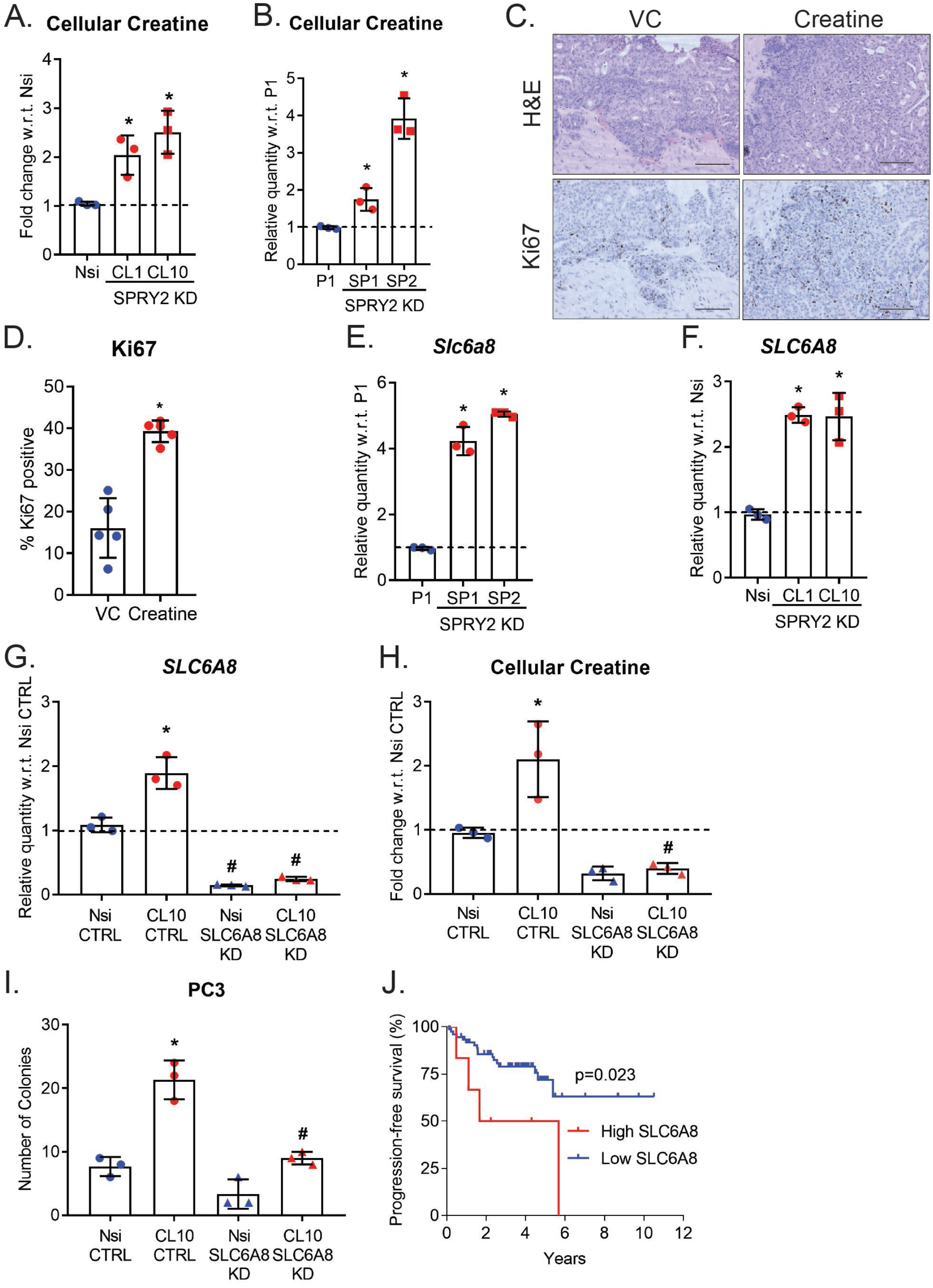
Increased cellular creatine sustains elevated growth rate of SPRY2 deficient cells. (A-B) Relative cellular creatine levels in human (A) and murine (B) prostate cancer cells (n=3 independent experiments; mean values ± SD are shown; *p<0.05 compared to Nsi and P1, respectively; unpaired t-test). (C) Representative H&E and Ki67 images of prostate tumour sections from *Pten*^*pc-/-*^ *Spry2*^*pc-/-*^ mice treated for two months with vehicle (VC) or 1% creatine. (n=5 mice for each group; scale bar = 100 μm). (D) IHC quantification of Ki67 in prostate tumours from *Pten*^*pc-/-*^ *Spry2*^*pc-/-*^ mice treated for two months with vehicle (VC) or 1% creatine (n=5 mice for each group; mean values ± SD are shown; *p<0.05; Mann-Whitney test). (E) Relative levels of creatine transporter, *Slc6a8* mRNA in murine prostate cancer cells (data shown are representative of n=3 independent experiments; mean values ± SD are shown; *p<0.05 compared to P1; 1-way ANOVA with Dunnett’s multiple comparison test). (F) Relative levels of creatine transporter, *SLC6A8* in human prostate cancer cells (n=3 independent experiments; mean values ± SD are shown; *p<0.05 compared to Nsi; 1-way ANOVA with Dunnett’s multiple comparison test). (G) Relative levels of *SLC6A8* mRNA in PC3 Nsi and SPRY2 KD CL10 with SLC6A8 KD (n=3 independent experiments; mean values ± SD are shown; *p<0.05 compared to Nsi CTRL & #p<0.05 compared to respective CTL cells; 1-way ANOVA with Tukey’s multiple comparison test). (H) Relative cellular creatine levels in PC3 Nsi and SPRY2 KD CL10 with SLC6A8 KD (n=3 independent experiments; mean values ± SD are shown; *p<0.05 compared to Nsi CTRL & #p<0.05 compared to respective CTL cells; 1-way ANOVA with Tukey’s multiple comparison test). (I) Number of soft agar colonies of PC3 Nsi and SPRY2 KD CL10 with SLC6A8 KD (data shown are representative of n=3 independent experiments; mean values ± SD are shown; *p<0.05 compared to Nsi CTRL & #p<0.05 compared to respective CTRL cells; 1-way ANOVA with Tukey’s multiple comparison test). (J) Kaplan–Meier plot for progression‐ free survival in MSKCC, Cancer Cell 2010 (Taylor *et al*., 2010) prostate cancer dataset showing cases with high (more than z-score= 1.8; n=6) or low (less than z-score= 1.8; n=74) expression of *SLC6A8*; log‐ rank Mantel-Cox test.

To investigate if elevated levels of creatine is causative for enhanced tumour growth, we tested the impact of creatine supplementation on tumour growth of *Pten* and *Spry2* double deficient mice. We treated six months old *Pten*^*pc-/-*^ *Spry2*^*pc-/-*^ mice with creatine for two months. Creatine treatment significantly increased the proliferation of prostate tumours with enhanced Ki67 staining (Figure 2C-D, EV2B). Furthermore, the creatine *SLC6A8* transporter was expressed at significantly higher levels in both human and murine prostate cancer cells deficient for SPRY2 (Figure 2E-F). Cellular creatine can be taken up via SLC6A8 as well as synthesised *de novo* from glycine and arginine (EV2A). To further characterise the source of cellular creatine, we stably knocked down SLC6A8 in SPRY2 deficient PC3 cells (Figure 2G). SLC6A8 deficiency significantly decreased cellular creatine levels in PC3 cells (Figure 2H). Functionally, SLC6A8 deficiency significantly decreased soft agar colony formation efficiency in SPRY2 deficient PC3 cells (Figure 2I), thus supporting the idea that creatine uptake is required for SPRY2 deficient driven growth. In the MSKCC clinical prostate cancer cohort (c-Bio portal) (Taylor *et al*, 2010), patients with high *SLC6A8* expression were at risk of disease progression (Figure 2J), and expression of *SPRY2* and *PTEN* negatively correlated with *SLC6A8* (EV2C-D). Collectively, SLC6A8 mediated creatine uptake supports the growth of PTEN and SPRY2 double deficient prostate cancer cells.

### Cyclocreatine suppresses the phosphagen system

Cellular creatine and creatine kinases are components of the phosphagen system (Ellington, 2001) which serves as a cellular energy buffering system to quickly generate ATP on demand (Abida *et al*., 2019). By efficiently storing high energy phosphates as phosphocreatine, the phosphagen system may promote tumorigenesis, especially in rapidly proliferating tumours with high energy demands. Since creatine can stimulate mitochondrial respiration in certain tissues (Kazak *et al*, 2015), we investigated the effect of creatine on cellular respiration. Creatine treatment significantly induced basal respiration in SPRY2 deficient PC3 cells (CL1 and CL10) (Figure EV2E-G). Consistent with creatine induced basal respiration, the creatine kinase activity (namely generation of phosphocreatine and ADP from creatine and ATP) was also elevated in SPRY2 deficient cells (Figure 3A). Despite increased creatine kinase activity and creatine induced basal respiration, the cellular ATP and phosphocreatine levels were not altered in SPRY2 deficient cells (Figure 3B-C).

**Figure 3.**
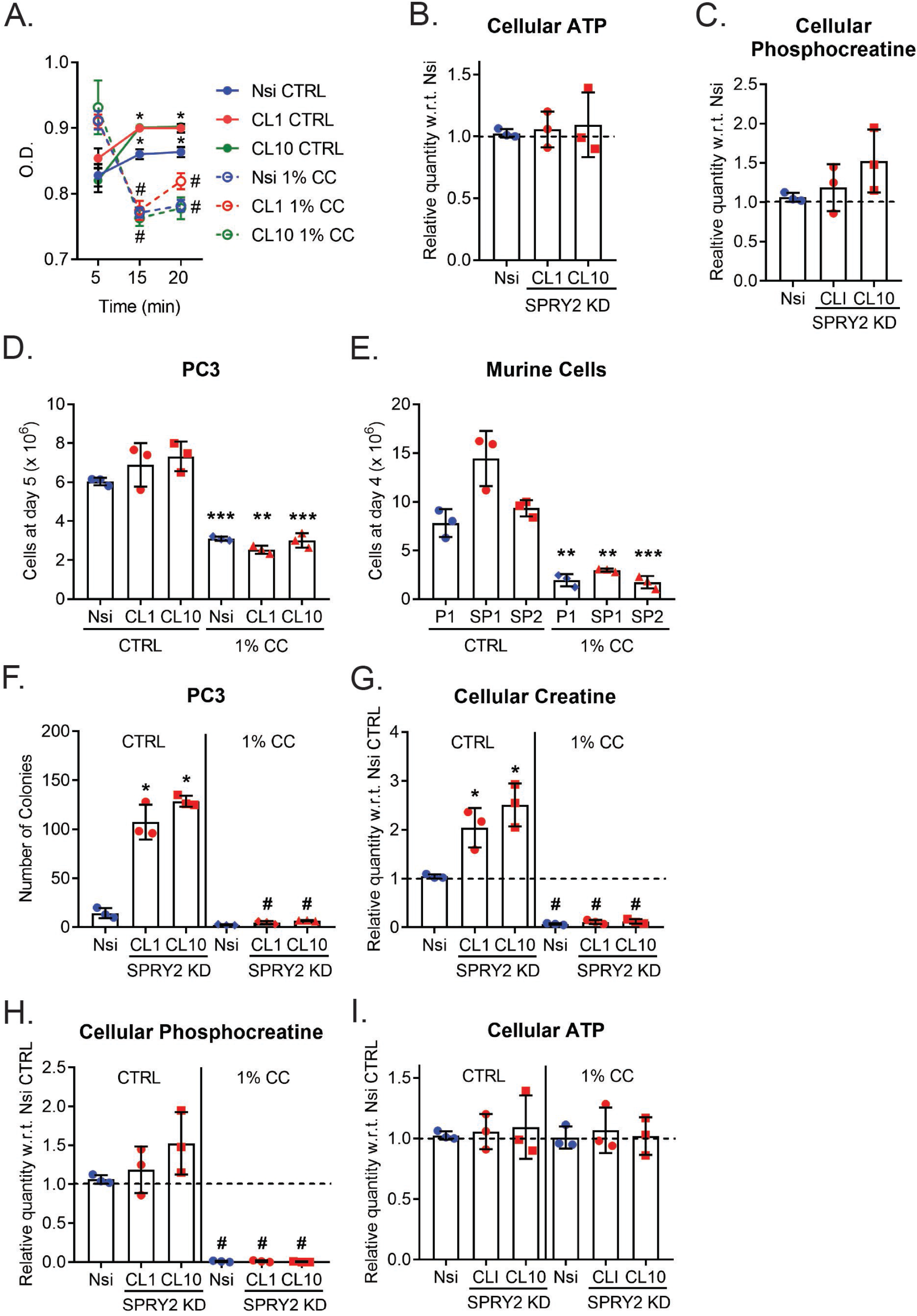
Phosphagen system in SPRY2 deficient cells. (A) Creatine kinase activity (conversion of creatine and ATP to phosphocreatine and ADP) in control and 1% cyclocreatine (CC) treated PC3 Nsi and SPRY2 KD clones CL1 and CL10 (n=3 independent experiments; mean values ± SD are shown; *p<0.05 compared to Nsi; #p<0.05 comparing 1% CC treated with untreated; 2-way ANOVA with Holm-Sidak’s multiple comparison test). (B-C) Relative levels of cellular ATP (B) and phosphocreatine (C) in PC3 Nsi and SPRY2 KD clones CL1 and CL10 (n=3 independent experiments; mean values ± SD are shown). (D-E) *In vitro* growth of human PC3 (E) and murine (F) cells treated with control (CTRL) or 1% cyclocreatine (CC) (n=3 independent experiments; mean values ± SD are shown; **p<0.005, ***p<0.0005 comparing CC treatment with respective CTRL cells; unpaired t-test). (F) Soft agar colony quantifications of PC3 Nsi and SPRY2 KD clones (CL1 and CL10) treated with control or 1% cyclocreatine (CC) containing medium (data shown are representative of n=2 independent experiments; mean values ± SD are shown; *p<0.05 compared to Nsi CTRL & #p<0.05 compared to respective CTRL cells; 1-way ANOVA with Tukey’s multiple comparison test). (G-I) Relative levels of cellular creatine (H), phosphocreatine (I) and ATP (J) in control (CTRL) or 1% cyclocreatine (CC) treated PC3 Nsi and SPRY2 KD clones CL1 and CL10 (n=3 independent experiments; mean values ± SD are shown; *p<0.05 compared to Nsi CTRL & #p<0.05 compared to respective CTL cells; 1-way ANOVA with Tukey’s multiple comparison test).

Cyclocreatine is a creatine analogue that blocks the phosphagen system by displacing creatine. We applied cyclocreatine treatment (1%) to investigate functional effects of creatine metabolism. Cyclocreatine suppressed *in vitro* creatine kinase activity (Figure 3A) and impaired *in vitro* proliferation of both human and murine prostate cancer cells (Figure 3D-E). Furthermore, cyclocreatine treatment significantly impaired the colony forming ability of PC3 cells (Figure 3F), along with reduced cellular creatine and phosphocreatine levels (Figure 3G-H) while the ATP levels were not altered (Figure 3I). Together these data suggest cyclocreatine potently suppresses activities of the phosphagen system.

### Creatine metabolism regulates cellular S-adenosyl methionine (SAM) levels

Consistent with the observation of enhanced creatine uptake in SPRY2 deficient cells, in creatine proficient serum supplemented culture conditions, cyclocreatine drastically reduced creatine uptake (Figure 4A). In addition, in serum-free culture conditions, we observed enhanced creatine synthesis in SPRY2 deficient PC3 cells (Figure 4A), which was accompanied by significantly higher levels of phosphocreatine (Figure 4B). Treatment with cyclocreatine significantly reduced cellular levels of creatine and phosphocreatine irrespective of SPRY2 status (Figure 4A-B), with both creatine uptake and biosynthesis suppressed. Collectively, cyclocreatine treatment blocked creatine uptake, creatine biosynthesis and phosphocreatine formation.

**Figure 4.**
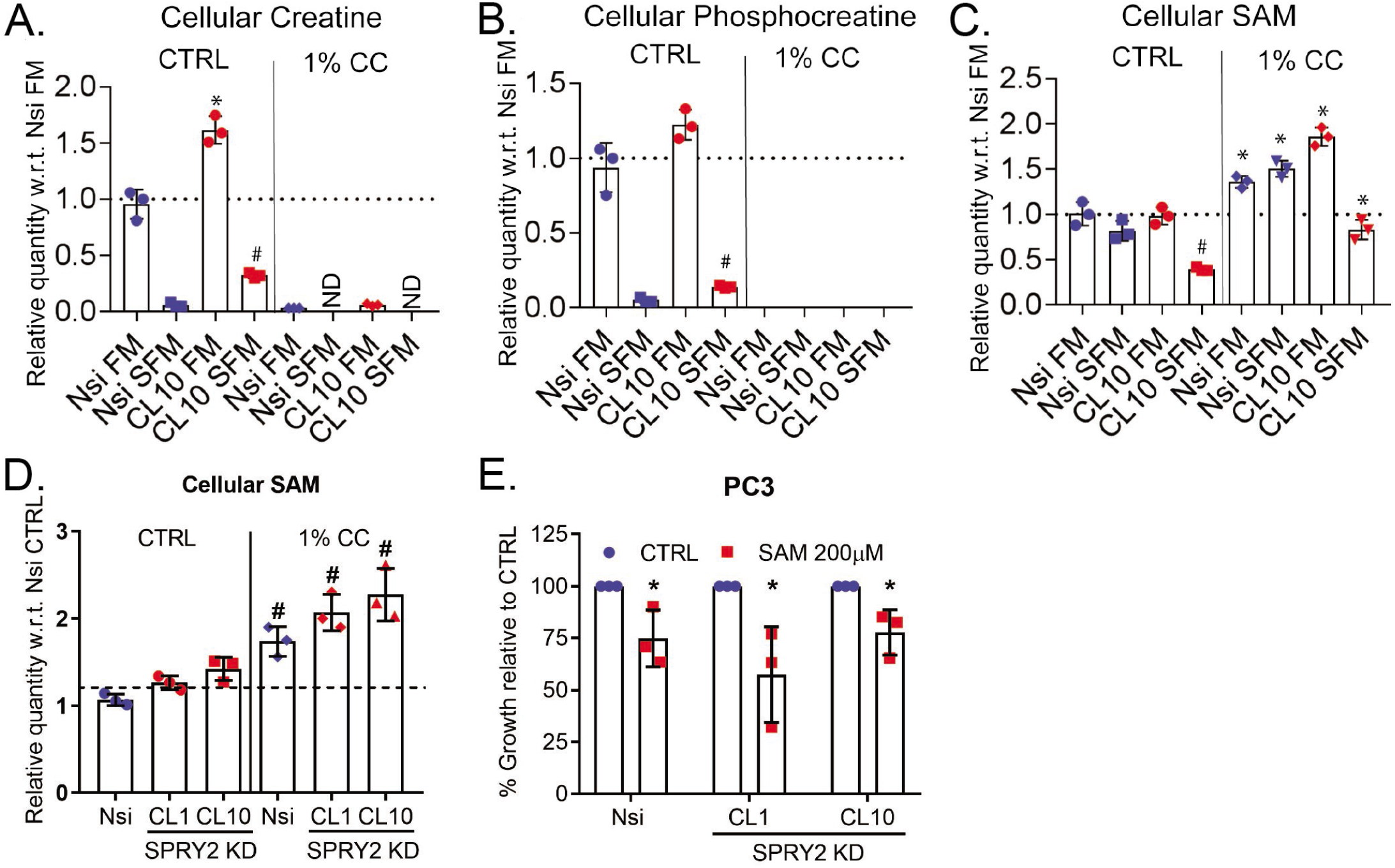
Cyclocreatine treatment induces growth arrest and SAM accumulation. (A-B) Relative levels of cellular creatine (A) and phosphocreatine (B) in PC3 Nsi and SPRY2 KD clone CL10 grown in either control or 1% cyclocreatine (CC) containing serum free medium (SFM) or medium with 10% serum (FM) (n=3 technical replicates; mean values ± SD are shown; *p<0.05 compared to Nsi CTRL & #p<0.05 compared to respective CTL cells; 2-way ANOVA with Tukey’s multiple comparison test). (C) Relative levels of cellular SAM in PC3 Nsi and SPRY2 KD clone CL10 grown in either control or 1% cyclocreatine (CC) containing serum free medium (SFM) or medium with 10% serum (FM) (n=3 technical replicates; mean values ± SD are shown; *p<0.05 compared to Nsi CTRL & #p<0.05 compared to respective CTL cells; 2-way ANOVA with Tukey’s multiple comparison test). (D) Relative levels of cellular S-adenosyl methionine (SAM) PC3 Nsi and SPRY2 KD clones (CL1 and CL10) treated as indicated (n=3 independent experiments; mean values ± SD are shown; #p<0.05 compared to respective CTL cells; 1-way ANOVA with Tukey’s multiple comparison test). (E) Growth of indicated cells in the presence of 200 µM S-adenosyl methionine (SAM) relative to respective cells treated with control medium (n=3 independent experiments; mean values ± SD are shown; *p<0.05 compared to respective CTRL cells; Multiple t-tests using Holm-Sidak method).

S-adenosyl methionine (SAM) is required for *de novo* creatine synthesis. In the presence of enhanced cellular creatine synthesis, cellular SAM steady state level may be reduced. We carried out targeted metabolomics analysis to study metabolic changes associated with SPRY2 deficiency with and without cyclocreatine treatment. In the absence of exogenous creatine, SPRY2 deficient cells (with enhanced creatine biosynthesis) exhibited significantly reduced levels of SAM (Figure 4C left side), consistent with increased SAM utilisation for creatine biosynthesis. We therefore reasoned that blockade of creatine biosynthesis may reciprocally lead to SAM accumulation. Indeed, cyclocreatine treatment significantly increased the cellular level of SAM in PC3 cells regardless of SPRY2 status and culture condition (Figure 4C right side, 4D). Consistent with a previous report (Schmidt *et al*, 2016), SAM treatment decreased the growth of PC3 cells independent of SPRY2 status (Figure 4E). Hence, in addition to the direct effects of reduced creatine levels, cyclocreatine induced effects may result in part from the accumulation of SAM (due to blockade of creatine biosynthesis).

### Effects on cyclocreatine treatment on *in vivo* prostate carcinogenesis

To investigate the tumour suppressive effects of cyclocreatine, we treated prostate tumour bearing *Pten*^*pc-/-*^ *Spry2*^*pc-/-*^ mice with 1% cyclocreatine for one month. Cyclocreatine treatment significantly decreased tumour cell proliferation with reduced Ki67 staining (Figure 5A-B), while the trend for reduced tumour weights did not reach statistical significance (Figure 5C). Regardless of the tumour genotype, the process of cancer metastasis is metabolically demanding for both energy and biomass. We therefore applied a liver metastasis model with splenic injection of PC3M cells to test the impact of manipulation of the SAM-creatine mediated metabolism by cyclocreatine in metastasis. Treatment with 1% cyclocreatine for a month drastically decreased liver metastatic burden of injected PC3M prostate cancer cells (Figure 5D-E).

**Figure 5.**
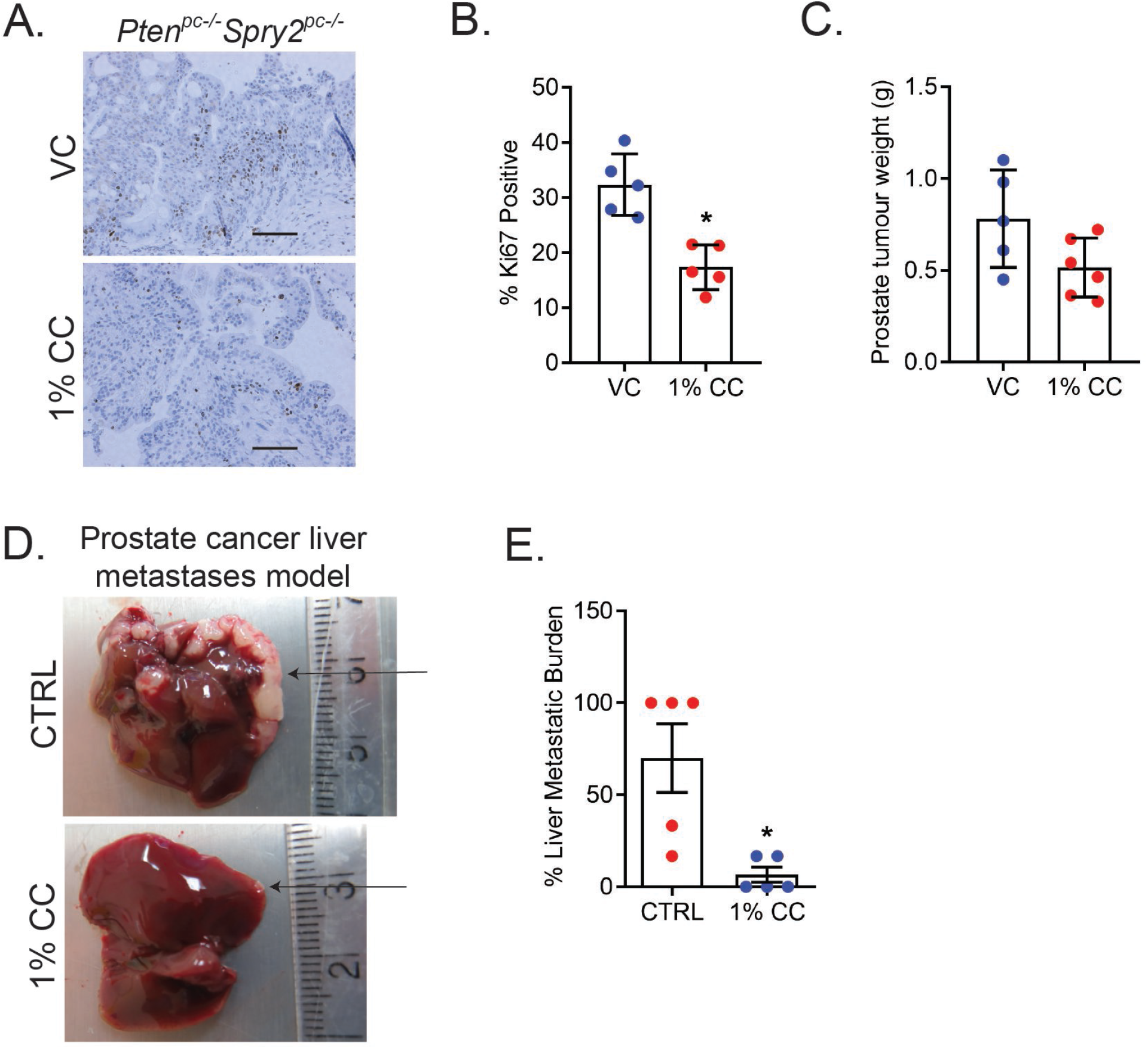
Cycocreatine treatment and *in vivo* prostate cancer progression. *(A)* Representative Ki67 images of prostate tumour sections from *Pten*^*pc-/-*^ *Spry2*^*pc-/-*^ mice treated for one month with vehicle (VC) or 1% cyclocreatine (n=5 mice for each group; scale bar = 100 μm). (B) IHC quantification of Ki67 in prostate tumours from *Pten*^*pc-/-*^ *Spry2*^*pc-/-*^ mice treated for two months with vehicle (VC) or 1% cyclocreatine (CC) (n=5 mice for each group mean values ± SD are shown; *p<0.05; Mann-Whitney test). (C) Non-cystic prostate tumour weights from *Pten*^*pc-/-*^ *Spry2*^*pc-/-*^ mice treated for one month with vehicle (VC; n=5) or 1% cyclocreatine (CC; n=6); mean values ± SD are shown). (D) Representative images of PC3M liver metastases in CD-1 nude mice treated with control or 1% cyclocreatine (CC) water (*ad libitum*) for one month (n=5 mice for each group; arrows indicate liver metastases). (E) Percentage PC3M liver metastases burden in CD-1 nude mice treated with control or 1% cyclocreatine (CC) water (*ad libitum*) for one month (n=5 mice for each group; mean values ± SEM are shown; *p<0.05; Mann-Whitney test).

Overall, our data highlighted the importance of creatine metabolism in prostate cancer and identified a potential role of SAM utilisation during creatine biosynthesis as an additional control of cancer growth (Figure 6).

**Figure 6.**
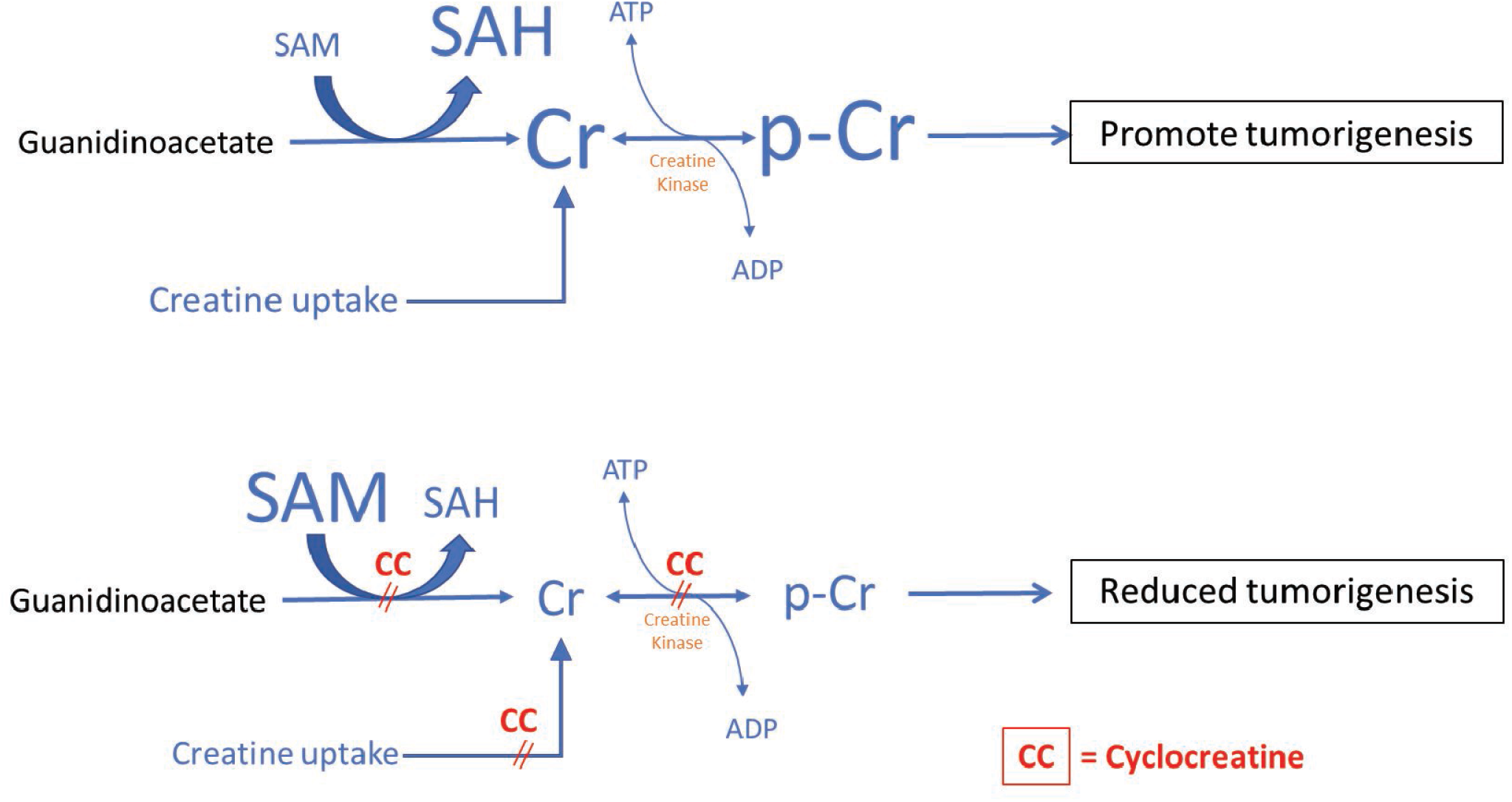
Schematic representation of creatine metabolism in prostate cancer and how cyclocreatine (CC) treatment suppresses prostate carcinogenesis.

## DISCUSSION

Metabolic adaptations in cancers (including prostate cancer) may contribute to progressive tumour growth. Understanding oncogenic metabolic alterations will highlight relevant tumour metabolic vulnerability to inform novel strategies to overcome cancer progression. Hence, we investigated the metabolic changes observed in prostate cancer driven by clinically-relevant deficiency of PTEN and SPRY2. Building on previous research (Patel *et al*., 2018; Patel *et al*., 2013), combined loss of the tumour suppressors PTEN and SPRY2 was found to further enhance tumour growth and progression. Here we show that creatine metabolism (uptake and biosynthesis) supports the growth of PTEN and SPRY2 deficient prostate cancer while cyclocreatine mediated blockade of the phosphagen system suppressed proliferation.

Diet is a major source of creatine. Circulating creatine can enter cells via the bi-directional SLC6A8 creatine transporter. PTEN and SPRY2 deficient cells have upregulated SLC6A8 expression to maintain elevated cellular creatine levels required for enhanced proliferation. In addition, combined PTEN and SPRY2 deficiency also promotes creatine biosynthesis to sustain cell growth. We find that PTEN and SPRY2 deficient cells are able to utilise exogenous creatine to stimulate basal respiration, and exhibit elevated creatine kinase activity.

We have previously demonstrated that PTEN and SPRY2 deficiency drives prostate cancer progression through enhanced HER2 signalling (Gao *et al*., 2012). HER2 mediated oncogenic signals can promote mitochondrial creatine kinase stability to enhance cellular bioenergetics (with increased phosphocreatine and cellular ATP levels) (Ben-Sahra & Puissant, 2018; Kurmi *et al*, 2018). Although we have not directly investigated the expression status of creatine kinases, we reason that the observed increased creatine kinase activity in PTEN and SPRY2 deficient prostate cancer cells is likely a result of altered expression of creatine kinases. However, we did not detect alterations in the overall levels of phosphocreatine and ATP in PTEN and SPRY2 deficient cells, which may reflect dynamic flux of the phosphagen system and tightly regulated nature of cellular energy homeostasis. Together and in support of recent reports (Kazak & Cohen, 2020; Kazak *et al*, 2019; Tang *et al*, 2015), our data is consistent with a functional role of creatine metabolism beyond energy buffering and ATP generation. Importantly, by blocking creatine uptake and biosynthesis using cyclocreatine, we identified SAM utilisation as a potential mechanism whereby changes in creatine metabolism may indirectly influence cancer growth. The functional and clinical significance of altered SAM metabolism warrants future investigations. Our data implicate cyclocreatine treatment in suppressing both creatine uptake and biosynthesis, although it remains to be clarified whether the observed effect on creatine biosynthesis represents a consequence of reduced creatine utilisation due to functional inhibition of creatine kinases.

The concept of resistance training is being considered as means to tackle ADT-induced loss of lean body mass. The idea of creatine supplement is studied as a component of ongoing lifestyle intervention clinical trial in prostate cancer (Fairman *et al*, 2019). Our observation of increased cellular proliferation in tumours from *Pten*^*pc-/-*^ *Spry2*^*pc-/-*^ mice following creatine supplement for only eight weeks highlights the need to assess creatine supplement in patients with caution.

Overall, we show that SPRY2 and PTEN double deficient prostate cancer cells sustain growth by engaging creatine metabolism to stimulate cellular respiration and regulate SAM levels. Furthermore, blockade of the phosphagen system produces anti-tumorigenic effects regardless of the status of SPRY2. Therapy such as cyclocreatine that targets the phosphagen system may bring about anti-tumour effects through suppressed creatine metabolism and accumulation of cellular SAM.

## METHODS

### Mice

All the animal experiments were carried out with ethical approval from the University of Glasgow under the revised Animal (Scientific Procedures) Act 1986 and the EU Directive 2010/63/EU (PPL P5EE22AEE). The ARR2Probasin-Cre (Pb-Cre), *Pten*^flox^, *Spry2*^flox^ (Patel *et al*., 2018; Patel *et al*., 2013) mice have been previously described, and applied to generate the *Pten*^*pc-/-*^ *Spry2*^*pc-/-*^experimental cohort. All mice in this study were on a mixed background and were genotyped by Transnetyx™ using PCR analysis of ear notch tissue. Mice were housed at 19°C to 23°C with a 12-hour light-dark cycle, and were fed a conventional diet with mains water ad libitum. They were housed in an enriched environment, with igloos, cardboard tubes and chewing sticks. Genotype-specific male mice were handled and aged until experimental time-points or ethically approved clinical endpoints (including palpable tumour with maximum diameter of 1.2 cm).

To investigate the effects of creatine supplement on GEMM prostate tumour growth, creatine (40 mg daily) (Sigma #C0780) was administered via gavage of 1% creatine solution for two months. Similarly, 1% cyclocreatine (w/v) (2-imino-1-imidazolidineaceticacid) (Sigma #377627) was supplied in drinking water *ad libitum* for one month.

CD-1 Nudes 6-8 weeks old male mice were used to study the extent of liver metastases following splenic injection of PC3M cells (five million cells per injection into the spleen of each experimental mouse). 1% cyclocreatine (w/v) (2-imino-1-imidazolidineaceticacid) (Sigma #377627) in drinking water was started one week after tumour cell implantation *ad libitum* for one month. The ARRIVE guidelines were used to design and execute experiments. Matastasis burden in the liver was obtained histologically as previously described (Patel *et al*., 2018).

### Cell Lines

Human PC3 and PC3M cells were purchased from ATCC. PC3M cells are metastasis-derived variant of human prostate cancer PC3 cells. Cell lines were mycoplasma negative and authenticated by LGC standards. P1 murine prostate cancer cells (RRID:CVCL_VQ82) were derived from a prostate tumour from a mouse bearing *Pten*^*pc-/-*^ tumour at 14 months. Similarly. SP1 and SP2 cell lines (RRID:CVCL_VQ84; RRID:CVCL_VQ85) were generated from two *Pten*^*pc-/-*^ *Spry2*^*pc+/-*^ tumour bearing endpoint mice at 6 and 8 months, respectively. Cell lines were routinely grown *in vitro* in media (RPMI for PC3 and PC3M and DMEM for P1, SP1 and SP2) supplemented with 10% FBS and 2 mM glutamine.

All siRNA and plasmid transfection experiments in PC3 cells were performed using nucleofection (Lonza-kit V)(see supplementary information for details). For stable SPRY2 KD, a 19-mer SPRY2 target sequence (5′-AACACCAATGAGTACACAGAG-3) (QIAGEN) was used and for the control a 19-mer NSi control sequence (QIAGEN) was used. Stable expression of ShSPRY2 and ShNSi in human PC3 cells was achieved using a plasmid pTER+ to insert the sequence of interest and selected using zeocin (300 μg/ml).

## Acknowledgements

This work was supported by Cancer Research UK (A17196 - core funding to the CRUK Beatson Institute, A15151 - awarded to HYL). LR received a CRUK Clinical Research Fellowship (A19493). We thank Arnaud Blomme for useful discussion.

## Author contributions

LR, RP, CAF, OJS and HYL designed research; RP, LR, CAF, JF, EM, TZ, DW, GM, and DS, performed research; LR, CAF, RP, LKR and HYL analysed and reviewed data; and LKR, CAF, RP and HYL wrote the paper.

## Expanded View Figure Legends

**Figure EV1. Concomitant PTEN and SPRY2 deficiencies in prostate tumours**

*(A)* Representative prostate tumour images from the indicated mice (n=7 mice for each group; scale bar = 1 cm).

(B) Western blot showing Sprouty2 expression in murine P1, SP1 and SP2 cells (data shown are representative of n=3 independent experiments; GAPDH is used a loading control).

(C) Western blot showing Sprouty2 expression in PC3 human prostate cancer cell lines as indicated human (data shown are representative of n=4 independent experiments; HSC70 is used as a loading control).

**Figure EV2. Creatine metabolism in prostate cancer**

*(A)* Schematic of creatine biosynthesis and metabolism: Creatine can be synthesised from glycine and arginine. Alternatively, creatine can be taken up by cells from circulation through the creatine transporter SLC6A8. Cellular creatine can be converted to phosphocreatine, which may be utilised as cellular energy buffer generating ATP upon demand.

(B) Non-cystic prostate tumour weights from *Pten*^*pc-/-*^ *Spry2*^*pc-/-*^ mice treated for two months with vehicle (VC; n=5) or 1% creatine (n=6) (mean values ± SD are shown).

(C-D) Correlation analyses of *SLC6A8* with *SPRY2* (C) and *PTEN* (D) mRNA in MSKCC, Cancer Cell 2010 (Taylor *et al*., 2010) prostate cancer dataset.

(E-G) Oxygen consumption rate in PC3 Nsi (E), SPRY2 KD CL1 (F) and SPRY2 KD CL10 (G) treated as indicated (n=4 independent experiments; mean values ± SEM are shown; *p<0.05 compared to CTRL; 2-way ANOVA with Tukey’s multiple comparison test).

